# Sparse dimensionality reduction for analyzing single-cell-resolved interactions

**DOI:** 10.1101/2024.12.01.626228

**Authors:** Niklas Brunn, Maren Hackenberg, Tanja Vogel, Harald Binder

## Abstract

**Summary:** Several approaches have been proposed to reconstruct interactions between groups of cells or individual cells from single-cell transcriptomics data, leveraging prior information about known ligand-receptor interactions. To enhance downstream analyses, we present an end-to-end dimensionality reduction workflow, specifically tailored for single-cell cell-cell interaction data. In particular, we demonstrate that sparse dimensionality reduction can pinpoint specific ligand-receptor interactions in relation to clusters of cell pairs. For sparse dimensionality reduction, we focus on the Boosting Autoencoder approach (BAE). Overall, we provide a comprehensive workflow, including result visualization, that simplifies the analysis of interaction patterns in cell pairs. This is supported by a Jupyter notebook that can readily be adapted to different datasets.

**Availability and implementation:** https://github.com/NiklasBrunn/Sparse-dimension-reduction

**Supplementary material:** …

## Introduction

Recent enhancements of techniques for measuring gene expression levels of cells along with multiple modalities such as their spatial localization in the tissue have led to a rapidly evolving research field for computational tools that enable a detailed study of cell-cell communication (Armingol et al. [2024]). Pioneering approaches integrate prior knowledge about known ligand-receptor interactions into a computational framework for reconstructing directed cell-cell interactions from transcriptomics data by leveraging the aggregated ligand expressions in sender cells and the aggregated cognate receptor expressions in receiver cells, of a specific type respectively (Efremova et al. [2020], Cabello-Aguilar et al. [2020], Jin et al. [2021]).

However, treating cells of the same type as a single entity may overlook potential within-cell type variability of interaction patterns (Armingol et al. [2024]). More recent approaches thus compute cell-cell interaction scores at the single-cell level (Li and Yang [2022], Raredon et al. [2023], Wilk et al. [2024], Bafna et al. [2023]). For example, NICHES^1^ (Raredon et al. [2023]) and Scriabin (Wilk et al. [2024]) construct cell-cell interaction matrices (CCIMs) based on known ligand-receptor interactions using single-cell gene expression data. Such a CCIM consists of the interaction scores for each active ligand-receptor interaction (features) in pairs of single cells (observations).

In the analysis of interaction patterns in cell pairs, the first step typically is dimensionality reduction, followed by clustering. Subsequent cluster-dependent differential interaction analysis through post-hoc statistical testing then identifies ligand-receptor interactions that are specifically active within clusters (Raredon et al. [2023]).

Here, we demonstrate a workflow that uses sparse dimensionality reduction to identify characteristic ligand-receptor interactions for clusters of cell pairs in an end-to-end manner, i.e. with ligand-receptor identification already integrated into dimensionality reduction. While sparse dimensionality reduction can potentially be obtained by several approaches (Boileau et al. [2020], Song et al. [2021]), we focus on the Boosting Autoencoder^2^ approach (BAE) (Hackenberg et al. [2024]), which we have recently introduced in the context of analyzing single-cell gene expression data. The BAE provides an interpretable representation of cell pairs in a low-dimensional latent space, where each latent dimension is linked to a sparse characterizing set of ligand-receptor interactions by model-selected weights.

However, different signs in the learned weights can make the interpretation challenging. We demonstrate how specifically the BAE approach can be easily adapted for addressing this by integrating a soft clustering component into the neural network architecture.

The learned representation of cell pairs, cluster memberships, and selected interactions for the different clusters can be used to generate easy-to-interpret two-dimensional visualizations using UMAP (McInnes et al. [2018]). Additionally, ranked lists of selected interactions per cluster, along with their importance scores can be extracted from the model output.

We provide Jupyter notebooks in our GitHub repository to analyze interaction patterns in single-cell pairs, which can be easily adapted for different datasets and to explore the functionality of the proposed BAE approach.

## The Boosting Autoencoder

The Boosting Autoencoder (BAE) is a deep learning approach for sparse and interpretable representation learning, originally designed for the analysis of single-cell RNA sequencing data (Hackenberg et al. [2024]), which we adapt here for the analysis of cell-cell interactions. A BAE uses an autoencoder architecture comprising two concatenated neural networks, the encoder and the decoder. In the present setting of cell-cell interactions, the encoder defines a parametric mapping from the feature space of ligand-receptor interactions to a low-dimensional latent space and the decoder vice versa. The objective of the optimization is to minimize a reconstruction loss measuring the deviation of the model reconstruction to the input under the constraint of reducing the dimensionality during the forward pass.

To enable a direct connection of interactions to latent dimensions, the BAE encoder consists of a single-layer linear neural network that is parameterized solely by a weight matrix. During training, the BAE iteratively updates the encoder weight matrix based on componentwise boosting (Tutz and Binder [2006, 2007]), a stepwise feature selection approach. Specifically, the negative gradients of the autoencoder reconstruction loss with respect to the latent representation are used to construct responses, to which linear models of the input interactions are fitted in a stepwise manner. In each iteration, only the weight corresponding to the specific interaction that most improves the current linear model fit is updated. For each dimension of the latent representation, the coefficients of this linear model represent the corresponding row of the encoder weight matrix. As the encoder weights corresponding to the model coefficients are initialized from zero, model training results in a sparse weight matrix. The componentwise boosting algorithm is embedded into a gradient-based optimization framework for jointly finding optimal encoder weights and decoder parameters. It offers the advantage of incorporating user-defined constraints into the feature selection step, e.g., to obtain disentangled latent dimensions (Hackenberg et al. [2024]).

In our adaptation of the BAE for analyzing single-cell-resolved interaction patterns, we use the BAE with the disentanglement constraint to learn low-dimensional representations of cell pairs in largely uncorrelated latent dimensions, each characterized by a specific small set of ligand-receptor interactions. During parameter optimization, the BAE iteratively links interactions to latent dimensions, where the corresponding encoder weights can have different signs. Consequently, each latent dimension can capture two distinct groups of cell pairs, represented by opposite signs.

To further enhance the interpretation of learned interaction patterns in latent dimensions, we introduce the split-softmax transformation after the encoder. Specifically, this transformation allows to split two different groups of cell pairs potentially represented in the same latent dimension, while keeping track of the selected characterizing interactions for each group. The split-softmax transformation first splits the activation in latent dimensions per cell pair into their positive and negative version, followed by a softmax transformation, to model associations of cell pairs to different latent dimensions. The split-softmax transformation allows for an equal contribution of positive and negative signals in the latent representation, whereas direct application of the softmax function would map negative signals towards zero, thus overlooking potentially meaningful signals. Each output dimension of the split-softmax transformation represents a cluster to which observations can be assigned with a certain probability. The number of clusters is thereby controlled by the user-defined number of latent dimensions for the dimensionality reduction. In contrast to frequently used step-by-step approaches, where dimensionality reduction is followed by clustering and subsequent differential interaction analysis, our BAE approach smoothly optimizes all model components in an end-to-end manner. An overview of the model architecture is provided in Fig. 1a.

**Fig. 1.**
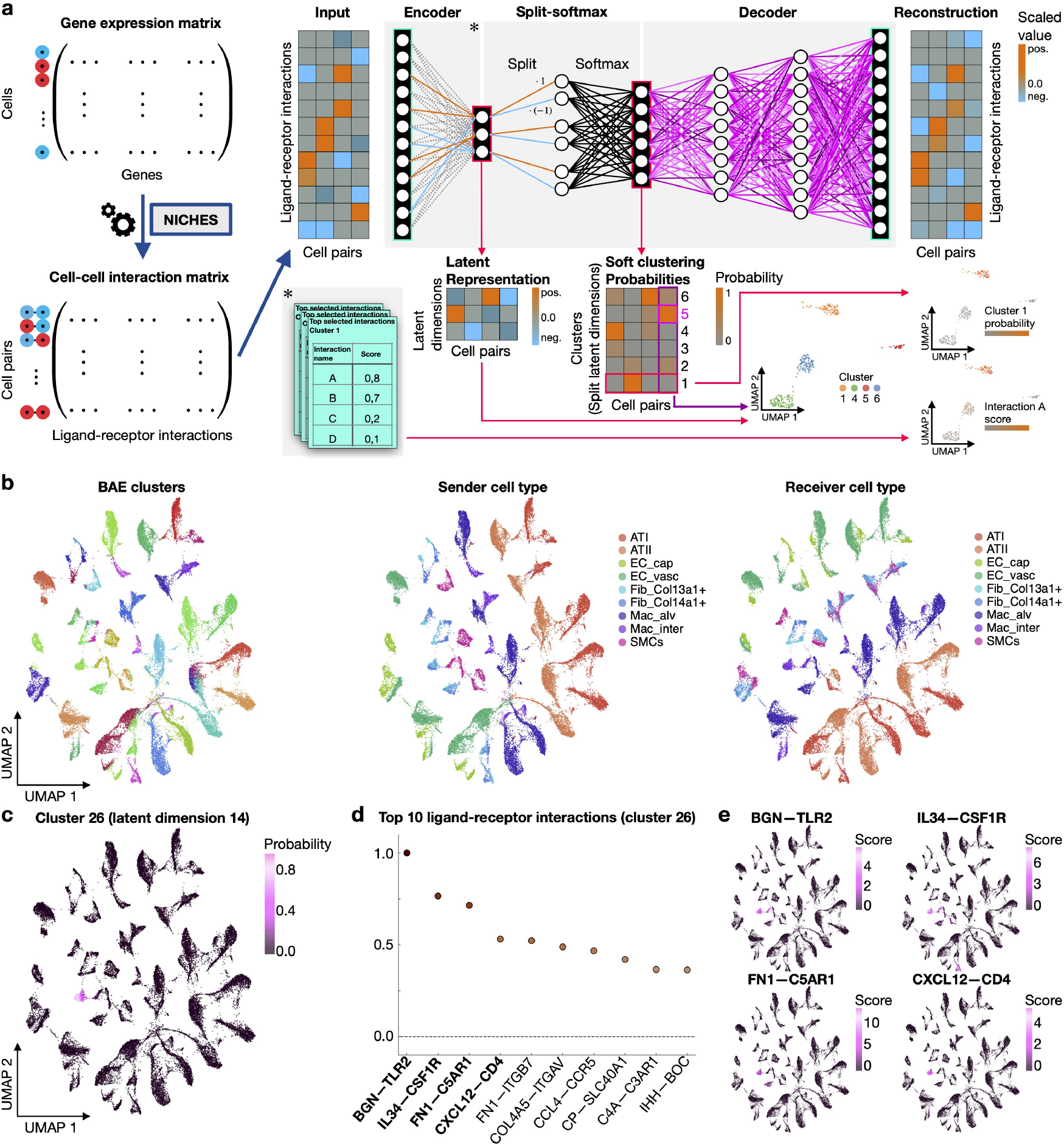
Schematic overview and application of the BAE. **a** (Left) The input data for training a BAE to learn single-cell interaction patterns is a CCIM constructed, e.g., by NICHES with standardized features. (Right) The BAE learns a structured and sparse connection of ligand-receptor interactions to latent dimensions (Encoder). The split-softmax transformation computes the cluster membership probabilities of cell pairs based on their representation. Each cell pair gets assigned to the cluster with the highest probability. The decoder network maps the split-softmax representation back into the ligand-receptor interaction space for reconstructing the interaction scores of cell pairs. 2D UMAP embeddings of the BAE latent representation can be used to visually inspect the results. Plots can be, e.g., colored by the cluster probabilities of cell pairs or scores of BAE-selected ligand-receptor interactions. **b-e** Result visualization of the BAE application to single-cell cell-cell interaction data computed by NICHES from rat data. The number of latent dimensions was set to *d* = 30, resulting in 60 clusters to which the model can assign cells. **b** (From left to right) 2D UMAP embedding of the latent representation of cell pairs colored by cluster membership. UMAP embedding of cell pairs colored by sender cell type. UMAP embedding of cell pairs colored by receiver cell type. **c** UMAP embedding colored exemplarily by the split-softmax representation of cell pairs in dimension 14 corresponding to cluster 26. **d** Scatter plot of the top selected ligand-receptor interactions for cluster 26. Dots represent min-max normalized positive encoder weights for the latent dimension 14 of the BAE in descending order (from left to right). **e** Top left/top right/bottom left/bottom right: UMAP embedding of cell pairs colored by interaction scores for the top 1/2/3/4 ligand-receptor interactions selected by the BAE for cluster 26.

## Analysis of single-cell interaction patterns

### Analysis workflow

In our proposed workflow for analysing single-cell-resolved interaction patterns, the input to a BAE is a CCIM, e.g., constructed by NICHES (Raredon et al. [2023]). NICHES can build a CCIM from either gene expression data from scRNA-seq or from a spatial transcriptomics experiment, ideally with single-cell resolution (Fig. 1a).

After training a BAE on the CCIM, it provides multiple outputs for exploring interaction patterns. First, a 2D UMAP representation of the data can be computed based on the learned latent representation, to visually explore similarities in the interaction patterns of cell pairs. Second, cell pairs are assigned to clusters with learned probabilities by the soft clustering component. Third, sparse lists of ranked ligand-receptor interactions help to pinpoint interactions that characterize clusters.

The BAE results can be visually inspected by coloring the 2D UMAP representation with the information from the different result modalities. Complementary metadata, such as cell type information, can thereby enhance the interpretation of the results.

Our implementation of the workflow can additionally handle spatial data and is accompanied by visualization functions that enhance the intuitive assessment of the learned latent patterns, making it easier for researchers to interpret and explore the data. Additionally, we provide Jupyter notebooks that allow users to either reproduce our analysis or to adapt it to other\ datasets, as well as to explore the functionality using simulated data.

### Visualization of results

As an examplary application, we considered a NICHES CCIM of scRNA-seq data from rat lungs (Raredon et al. [2019]).

We colored cell pairs in the 2D UMAP representation based on their cluster identity with the highest probability, the previously defined sender cell types, and the receiver cell types respectively (Fig. 1b). Comparing the three plots reveals that cell pairs with similar sender and receiver types have similar interaction profiles.

The individual clusters can be further examined for their specific characteristic interaction profiles (Fig. 1c-e). For example, inspection of the cluster membership probabilities of cell pairs in cluster 26 shows a separation from other cell pairs based on the latent patterns (Fig. 1c). The top characterizing interactions, which were selected by the BAE for cluster 26, are shown in Fig. 1d. These ligand-receptor interactions can be used to color the cell pair representation in the UMAP plot to see how well the interaction score patterns match the cluster identity of the cell pairs (Fig. 1e). Selected interactions of interest can be a basis for designing further laboratory experiments to confirm cluster-specific signaling.

## Conclusion

We presented a comprehensive workflow for sparse dimensionality reduction of single-cell-resolved interaction data, facilitated by the Boosting Autoencoder (BAE). To enhance the interpretability of the resulting low-dimensional embedding, we integrated a soft clustering component into the end-to-end approach. This integration enables the model to pinpoint specific interactions in relation to clusters of cell pairs, facilitating deeper insights into the underlying biological processes. With our workflow implementation, researchers can perform the analysis in a customizable Jupyter notebook with their own data and visualize the results.

## Data availability

The data underlying this article are available at https://zenodo.org/records/6846618 and can be processed for the analysis following the instructions and scripts in our GitHub repository https://github.com/NiklasBrunn/Sparse-dimension-reduction.

## Competing interests

No competing interest is declared.

## Funding

This work has been funded by the Deutsche Forschungsgemeinschaft (DFG, German Research Foundation) – Project-ID 322977937 – GRK 2344 (H.B, M.H., N.B., and T.V.) and Project-ID 499552394 – SFB 1597 (H.B. and M.H.).

## Author contributions statement

All authors contributed to the conception and design of the work. N.B. implemented the approach, and performed the computational analysis. H.B., M.H., and N.B. wrote the manuscript. T.V. commented on the manuscript. All authors approved the final version of the manuscript.

## Footnotes

1 https://github.com/msraredon/NICHES

2 https://github.com/NiklasBrunn/BoostingAutoencoder

## Notes

### Competing Interest Statement

The authors have declared no competing interest.

## References

E. Armingol, H. M. Baghdassarian, and N. E. Lewis. The diversification of methods for studying cell–cell interactions and communication. Nature Reviews Genetics, pages 1–20, 2024.

M. Bafna, H. Li, and X. Zhang. Clarify: cell–cell interaction and gene regulatory network refinement from spatially resolved transcriptomics. Bioinformatics, 39(Supplement 1):i484– i493, 2023.

P. Boileau, N. S. Hejazi, and S. Dudoit. Exploring high-dimensional biological data with sparse contrastive principal component analysis. Bioinformatics, 36(11):3422–3430, 2020.

S. Cabello-Aguilar, M. Alame, F. Kon-Sun-Tack, C. Fau, M. Lacroix, and J. Colinge. Singlecellsignalr: inference of intercellular networks from single-cell transcriptomics. Nucleic acids research, 48(10):e55–e55, 2020.

M. Efremova, M. Vento-Tormo, S. A. Teichmann, and R. Vento-Tormo. Cellphonedb: inferring cell–cell communication from combined expression of multi-subunit ligand–receptor complexes. Nature protocols, 15(4):1484–1506, 2020.

M. Hackenberg, N. Brunn, T. Vogel, and H. Binder. Infusing structural assumptions into dimension reduction for single-cell rna sequencing data to identify small gene sets. bioRxiv, pages 2024–02, 2024.

S. Jin, C. F. Guerrero-Juarez, L. Zhang, I. Chang, R. Ramos, C.-H. Kuan, P. Myung, M. V. Plikus, and Q. Nie. Inference and analysis of cell-cell communication using cellchat. Nature communications, 12(1):1088, 2021.

R. Li and X. Yang. De novo reconstruction of cell interaction landscapes from single-cell spatial transcriptome data with deeplinc. Genome biology, 23(1):124, 2022.

McInnes, J. Healy, and J. Melville. Umap: Uniform manifold approximation and projection for dimension reduction. arXiv preprint 1802.03426, 2018.

S. B. Raredon, T. S. Adams, Y. Suhail, J. C. Schupp, S. Poli, N. Neumark, K. L. Leiby, A. M. Greaney, Y. Yuan, C. Horien, et al. Single-cell connectomic analysis of adult mammalian lungs. Science advances, 5(12):eaaw3851, 2019.

M. S. B. Raredon, J. Yang, N. Kothapalli, W. Lewis, N. Kaminski, L. E. Niklason, and Y. Kluger. Comprehensive visualization of cell–cell interactions in single-cell and spatial transcriptomics with niches. Bioinformatics, 39(1):btac775, 2023.

D. Song, K. Li, Z. Hemminger, R. Wollman, and J. J. Li. scpnmf: sparse gene encoding of single cells to facilitate gene selection for targeted gene profiling. Bioinformatics, 37 (Supplement 1):i358–i366, 2021.

G. Tutz and H. Binder. Generalized additive modeling with implicit variable selection by likelihood-based boosting. Biometrics, 62(4):961–971, 2006.

G. Tutz and H. Binder. Boosting ridge regression. Computational Statistics & Data Analysis, 51(12):6044– 6059, 2007.

A. J. Wilk, A. K. Shalek, S. Holmes, and C. A. Blish. Comparative analysis of cell–cell communication at single-cell resolution. Nature Biotechnology, 42(3):470–483, 2024.

