## Supplementary Material for "Sparse dimensionality reduction for analyzing single-cell-resolved interactions"

**S2 NICHES** **4**

**S3 References** **5**

### S1 Boosting autoencoder

#### S1.1 Boosting autoencoder optimization algorithm

Compared to the original publication of the BAE [1], we optimized the implementation of the approach to speed up the computations. First, we applied weight decay to the decoder parameters as proposed in [2]. Regularizing the decoder parameters during optimization aids the model in retaining a sparse encoder weight matrix during training even if the model is trained for a larger number of epochs and prevents it from overfitting. Second, we now compute the gradients required for updating the parameters of all model components simultaneously, thereby speeding up the optimization process. Third, instead of only selecting and updating the optimal encoder weight for each latent dimension in each componentwise boosting iteration, we now allow all previously selected weights to be updated together with the current optimal weight. This provides more flexibility to the boosting model, which attempts to iteratively fit changing responses based on the negative gradients of the autoencoder reconstruction loss w.r.t. the latent representation throughout the interaction process. Finally, a soft clustering component can be integrated into the modeling framework by adding a $\mathrm{split\text{-}softmax}$ transformation after the encoder, as explained in the main text. Formally, for $n$ observations, let $\boldsymbol{Z} \in \mathbb{R}^{d\times n}$ denote the output of the encoder which is constructed by multiplying the data matrix $\boldsymbol{X} \in \mathbb{R}^{p\times n}$ with the sparse encoder weight matrix $\boldsymbol{W} \in \mathbb{R}^{d\times p}$, where $d \ll p$. The $\mathrm{split\text{-}softmax}$ transformation first splits the activation in latent dimensions per cell pair into their positive and negative version, followed by a $\mathrm{softmax}$ transformation $\sigma : \mathbb{R}^d \to \mathbb{R}^d$, to model associations of cell pairs to different latent dimensions. This can be formalized by the mapping

$$\begin{aligned} \sigma_{\text{split}} &: \mathbb{R}^d \to \mathbb{R}^{2d} \\ (z_1, \dots, z_d)^\top &\mapsto \sigma((z_1, -z_1, \dots, z_d, -z_d)^\top). \end{aligned}$$

#### S1.2 Boosting autoencoder architecture

We used the following BAE architecture for analyzing the CCIM constructed from the rat lung scRNA-seq data [3] by applying NICHES [4]. The encoder was defined to consist of one linear layer $f_{\text{enc}} : \mathbb{R}^p \to \mathbb{R}^d$, i.e., without a bias vector. The decoder consists of three layers of which two have trainable parameters. The first layer of the decoder is the $\mathrm{split\text{-}softmax}$ transformation $\sigma_{\text{split}} : \mathbb{R}^d \to \mathbb{R}^{2d}$ without trainable parameters, followed by a fully connected layer with $\tanh$ activation $f_{\text{dec}_1} : \mathbb{R}^{2d} \to \mathbb{R}^p$. The last layer was defined to be a fully connected affine layer $f_{\text{dec}_2} : \mathbb{R}^p \to \mathbb{R}^p$.

For the CCIM computed by NICHES using the subset of the rat lung data [3], the number of ligand-receptor interactions detected by NICHES was $p = 1740$ and for the BAE analysis we set $d = 30$, such that cell pairs could be assigned to $60$ different clusters. An exemplary model overview figure is provided in the main text in Fig. 1. a.

#### S1.3 Boosting autoencoder training

Prior to training, the BAE works best if the input features are standardized across the observations, i.e., z-scores of each feature vector were computed and used as the input for training the model. Each feature vector can either correspond to a detected ligand-receptor interaction by NICHES, consisting of the interaction scores of individual cell pairs, or to gene expressions of cells in a gene when analyzing single-cell gene expression data.

For training the model, decoder parameters were randomly initialized using the Xavier initialization strategy [5] and the encoder weights were set to zero for learning sparse connections.

We choose the mean squared error as the model reconstruction loss since the input data was standardized and used a well known modification of the Adam optimizer [6], AdamW [2], for optimizing the decoder parameters. The AdamW optimizer makes $L2$ regularization equivalent to weight decay, which is not the case for the vanilla Adam optimizer when adding a $L2$ penalty term for the decoder parameters to the reconstruction loss.

The BAE was trained for $2000$ epochs, where in each epoch, the input data was randomly divided into mini-batches of size $2^{12}$ and each mini-batch was used to update the parameters during each epoch.

The BAE optimization framework, consisting of a boosting component and the AdamW optimizer, makes use of gradient feedback, which is computed using the forward pass of the model with the current state of all model parameters using one mini-batch respectively. Negative gradients of the model loss with respect to the decoder parameters were then further passed to the AdamW optimizer, whereas negative gradients with respect to the latent representation of samples in the mini-batch in different latent dimensions were handed over to the componentwise boosting component for sparse feature selection and disentanglement of latent dimensions [1].

We set the learning rate for the AdamW optimizer to $0.01$, the weight decay parameter to $0.1$, the decay parameter for the first and the second momentum estimate to the default values, and the step size for the boosting component to $0.001$. We limited the number of boosting steps performed during each parameter update iteration for each latent dimension to $1$ since componentwise boosting is integrated into an iterative optimization scheme. More generally, we found that these values depict suitable candidates for default values for training a BAE, except for the batch size and the number of training epochs, which can be chosen based on the number of observations in the data and control the sparsity level of the resulting encoder weight matrix.

#### S1.4 Boosting autoencoder implementation

The BAE was implemented using the Julia programming language [7], v1.9.3 (macOS, aarch64). We mainly used the Flux (v0.14.15) library for deep learning models [8] which is based on the Zygote framework [9] for automatic differentiation. Other Julia packages required for the implementation are CSV v0.10.14, Clustering v0.15.7, ColorSchemes v3.25.0, DataFrames v1.6.1, Distances v0.10.11, IJulia v1.25.0, Plots v1.40.4, ProgressMeter v1.10.0, RCall v0.14.1, StatsBase v0.34.3, UMAP v0.1.11, VegaLite v3.3.0. All experiments were conducted on a MacBook Pro (2023) with an Apple M2 Max chip (12-core CPU, 38-core GPU) and 96 GB of RAM.

#### S1.5 Boosting autoencoder functionality

To demonstrate the functionality of the BAE with the disentanglement constraint, especially to highlight the capabilities of the soft clustering component, we added a tutorial notebook in our GitHub repository https://github.com/NiklasBrunn/Sparse-dimension-reduction. In this tutorial notebook users can run an exemplary BAE analysis on simulated binary scRNA-seq count data oriented on the simulation design proposed in [10]. In this case, the observations correspond to cells and the features correspond to genes.

#### S2 NICHES

NICHES (Niche Interactions and Communication Heterogeneity in Extracellular Signaling) is a recently published computational tool to reconstruct extracellular signaling at the single-cell level [3]. The NICHES software is implemented in the R programming language and is publicly available at https://github.com/msraredon/NICHES.

The algorithm takes as the input a normalized count matrix either resulting from scRNA-seq data or spatial transcriptomics (ST) data. In the latter case, NICHES makes use of information about the spatial locations of cells as an additional input to constrain the communication edges.

In addition, NICHES leverages prior knowledge about known ligand-receptor interactions from the FANTOM5 [11] or OmniPath database [12] for constructing a CCIM from the gene expression data.

Since consideration of each possible cell pair is costly both in computation time and memory, NICHES limits the computation of interaction scores to only a subset of the cell pairs. More precisely, when applied to a countmatrix of a scRNA-seq experiment, NICHES requires pre-annotated cell type information based on which cell pairs are uniquely sampled from. On the other hand, when spatial information is available, NICHES focuses on spatial neighbourhoods, i.e., niches.

NICHES can construct three different types of CCIMs: 1) A cell-to-cell interaction matrix, where observations correspond to cell pairs, 2) A cell-to-System matrix, where for each cell treated as a sender cell, the expression of the receptors are aggregated by either computing the sum or the mean of the receptor expressions in the receiver system. 3) A System-to-cell matrix, where for each cell treated as a receiver cell, the expressions of the ligands are aggregated by either computing the sum or the mean of the receptor expressions in the receiver system. Note that a system can have two different meanings depending on whether scRNA-seq data or ST data is used. Given scRNA-seq data, a system of a cell is defined by all cells that are connected to the cell after sub-sampling. For ST data, the system, or neighborhood, i.e., niche of a cell is defined by cells that are spatially close to the cell. Note that all types of CCIMs can be analyzed with the BAE.

For each individual cell pair and each ligand-receptor interaction, where ligands or receptors can consist of multiple components, NICHES computes an interaction score based on the product of the ligand and the receptor expression. In case of ligands or receptors consisting of multiple components, the final ligand or receptor expression that is used for the product for calculating the interaction score is defined by the geometrical mean of the individual expressions of the components. In case of a cell-to-system or a system-to-cell matrix, NICHES uses the aggregated ligand or receptor expressions within the systems for computing the interaction scores.

To handle high sparsity levels in the original gene expression matrix, the authors of NICHES recommend using an imputation algorithm, e.g., ALRA [13] prior to applying the NICHES algorithm. Note, that data imputation imputes gene expression values of genes whose expression levels were measured as zeros, where it is not clear weather those imputed values were hidden during measurement or if they are false positives.

#### S3 References

1. Hackenberg M, Brunn N, Vogel T, and Binder H. Infusing structural assumptions into dimension reduction for single-cell RNA sequencing data to identify small gene sets. bioRxiv 2024:2024–2.
2. Loshchilov I and Hutter F. Decoupled weight decay regularization. arXiv preprint arXiv:1711.05101 2017.
3. Raredon MSB, Adams TS, Suhail Y, Schupp JC, Poli S, Neumark N, Leiby KL, Greaney AM, Yuan Y, Horien C, et al. Single-cell connectomic analysis of adult mammalian lungs. Science advances 2019;5:eaaw3851.
4. Raredon MSB, Yang J, Kothapalli N, Lewis W, Kaminski N, Niklason LE, and Kluger Y. Comprehensive visualization of cell–cell interactions in single-cell and spatial transcriptomics with NICHES. Bioinformatics 2023;39:btac775.
5. Glorot X and Bengio Y. Understanding the difficulty of training deep feedforward neural networks. In: *Proceedings of the thirteenth international conference on artificial intelligence and statistics*. JMLR Workshop and Conference Proceedings. 2010:249–56.
6. Kingma DP and Ba J. Adam: A method for stochastic optimization. arXiv preprint arXiv:1412.6980 2014.
7. Bezanson J, Edelman A, Karpinski S, and Shah VB. Julia: A Fresh Approach to Numerical Computing. SIAM Review 2017;59:65–98.
8. Innes M. Flux: Elegant machine learning with Julia. Journal of Open Source Software 2018;3:602.
9. Innes M, Edelman A, Fischer K, Rackauckas C, Saba E, Shah VB, and Tebbutt W. A differentiable programming system to bridge machine learning and scientific computing. arXiv preprint arXiv:1907.07587 2019.
10. Hess M, Hackenberg M, and Binder H. Exploring generative deep learning for omics data using log-linear models. Bioinformatics 2020;36:5045–53.
11. Ramilowski JA, Goldberg T, Harshbarger J, Kloppmann E, Lizio M, Satagopam VP, Itoh M, Kawaji H, Carninci P, Rost B, et al. A draft network of ligand–receptor-mediated multicellular signalling in human. Nature communications 2015;6:7866.
12. Türei D, Valdeolivas A, Gul L, Palacio-Escat N, Klein M, Ivanova O, Ölbei M, Gábor A, Theis F, Módos D, et al. Integrated intra-and intercellular signaling knowledge for multicellular omics analysis. Molecular systems biology 2021;17:e9923.
13. Linderman GC, Zhao J, Roulis M, Bielecki P, Flavell RA, Nadler B, and Kluger Y. Zero-preserving imputation of single-cell RNA-seq data. Nature communications 2022;13:192.
